# Stability and characterization of infectious monkeypox virus extracellular virions in vitro and vivo; implications for transmission, pathogenesis and treatment

**DOI:** 10.1101/2025.03.12.642804

**Authors:** K. Ostrowski, C. Davis, S. Bixler, A. Garrison, Joseph W. Golden, A. Piper, D. J. Koehler, S. K. Ricks, Janice Williams, E.M. Mucker

**Affiliations:** Molecular Virology Branch, Virology Division, United States Army Medical Research Institute of Infectious Diseases; Applied Diagnostics Branch, Diagnostic Systems Division, United States Army Medical Research Institute of Infectious Diseases; Viral Pathogenesis Branch, Virology Division, United States Army Medical Research Institute of Infectious Diseases; Diagnostic Systems Division, United States Army Medical Research Institute of Infectious Diseases; Pathology Division, United States Army Medical Research Institute of Infectious Diseases

**Keywords:** extracellular, released, enveloped, cell associated virions, monkeypox, smallpox, variola, vaccinia, pathogenesis, stability, EEV, CEV, rCEV

## Abstract

Poxviruses exist as multiple infectious morphogenic forms commonly simplified as mature virions (MV) and extracellular virions (EV). The roles of morphogenic subtypes as related to disease and transmission are enigmatic as EVs can exist both as cell associated (CEV) or released particles (rCEV) each with potentially unique biochemical properties impacting stability and infectivity. *In vitro* analysis of prototypical poxviruses is commonly utilized to infer larger conclusions about the *in vivo* function of all EV-like particles. Here we show that infectious EV of MPXV and VACV strains are stable for ≥ 14 weeks and are more sensitive to human or non-human primate complement compared to rabbit-derived complement. We also characterize the levels of EV produced during MPXV infection in NHPs and found temporal differences in production that may influence spread. Also, we present data characterizing and contrasting the EV from monkeypox (MPXV) and vaccinia virus (VACV) strains. Specifically, we quantified infectious EV quantities produced by different cultured cell lines and characterized the infectious properties and composition of the released (extracellular) virions. We conclude that A33 neutralizable cell associated-like virions (CEV-like), a form of EV, can significantly increase, depending on the strain of VACV or MPXV and the cell lines from which they were released. Based on the outcomes of our studies, the importance of understanding specific orthopoxvirus EV roles in a host-specific manner, as it relates to pathogenesis, stability, and transmission, is warranted and requires further study.

## Introduction

Within the family *Poxviridae* are double stranded DNA-containing viruses that have an eclectic array of vertebrate and arthropod hosts(1). Within the family of complex viruses, many are associated with human disease and are mainly grouped within the *Orthopoxvirus* genus. The causative agents of smallpox, variola virus (VARV), and Mpox, monkeypox virus(MPXV), as well as the vaccine primarily utilized to eradicate smallpox disease, vaccinia virus (VACV) are a few examples.

Eradication of smallpox was only possible because of the human-restricted nature of VarV, but its known virulence and enigmatic mechanisms of diseases can only be assessed in two global locations and is limited to cell culture with few animal modeling systems(2, 3). The restriction of VarV to both use and location is highly regulated by the World Health Organization, further limiting the extent of our specific understanding of VarV. To fill in knowledge gaps about biology, virology, and human disease of smallpox, surrogate orthopoxviruses and their diseases have historically been utilized (e.g.,(4, 5)). Since MPXV causes a smallpox-like disease in humans, is zoonotic in nature, and has been endemically circulating in parts of Africa, it has been historically used as primarily surrogate of smallpox disease(5). Vaccinia virus is even more accessible and its importance as a human vaccine gives good justification for its use for both a general understanding of the virology of poxviruses and to diseases with orthopoxviral-related etiology(6–8).

Clinical presentation of MPXV in humans include fever and vesiculopustular rash that can rarely lead to systemic shock and death. Two clades of MPXV exist: Clade II found in West Africa(9–13) and Clade I from Africa’s Congo Basin(14–22). While there are reports of occasional travel and trade related cases of MPXV infection(23, 24), outbreaks were typically contained within Africa until the global emergence of Clade II MPXV in 2022(25, 26). The sustained circulation of Clade I MPXV in the current DRC outbreak (27, 28) resulted further spread to other countries including Kenya, Burundi, Nigeria, Thailand, Sweden, and the Philippines(29). On 14 August 2024, the World Health Organization (WHO) declared a Public Health Emergency of International Concern. Sequencing analysis of the outbreak identified a subclade of MPXV (Ib) that has several virulence gene deletions that complicate diagnostics with existing real-time PCR assays(27, 28, 30).

Given the pandemic potential of MPXV, the emergence of other poxviruses such as Alaskapox virus (31), and concern about reintroduction of VARV, further studies on the biology of poxvirus replication and transmission are needed. Poxviruses have a complex viral replication strategy that involves at least three antigenically distinct particles enveloped virus (EEV)(32, 33),. After initial assembly and processing of both viral and host proteins, a membrane derived enveloped particle is the first infectious form of the virus, the intracellular mature virion (IMV or MV). Select particles are then further wrapped with two host-derived membranes, intracellular enveloped virions (IEV), that are either immediately released (extracellular enveloped virions or EEV) or stay associated with external host cell membrane (cell associated enveloped virions or CEV) after fusion to endosomes or the host membrane itself (34). This results in an IMV-like particle with additional host cell membranes and are commonly discussed together as extracellular virions or EV.

### Terminology guide for manuscript

| Virion | Acronym | Summary Description | References |
| --- | --- | --- | --- |
| **MV** | mature virion | intracellular particles, typically synonymous with IMV; further differentiated by IMV-specific neutralizing antibodies | (1, 35) |
| **IMV** | intracellular mature virion | morphogenic first infectious form of poxvirus; mainly remains intracellular until cell lysis or may gain another cellular derived membrane for cell association and/or extracellular release; | (1, 34) (36) |
| **IMV-like** | intracellular mature virion-like | infectious intracellular virions with antigenic properties synonymous with IMV; usually determined by lack of addition envelopment; may contain particles that are more likely to be further processed for extracellular release; further differentiated by neutralization by IMV-specific antibodies | (37) |
| **EV** | Extracellular virion | any released particle detected in supernatant; typically, though synonymous with EEV/rCEV, but may likely contains MV and MV-like particles | (1, 35) |
| **CEV** | cell associated virions | Infectious morphogenic form of poxvirus virion that is retained on cell surface, thought to be responsible for rapid near-cell infection; | (34, 38) |
| **rCEV** | released cell associated virions | Infectious; cell associated virions that are released or are free of the cell and present in the supernatant after infection; enveloped and thought to be antigenically and functionally similar EEV; can also be released from the cell surface chemically (e.g., trypsin) or via antibodies (e.g., Mab c6C); defined sensitivity to anti-EEV antibodies, resistance to anti-MV neutralization and lack of neutralization by Mab c6C | (39) |
| **CEV-like** | Cell associated virion | Infectious; CEV and/or rCEV |  |
| **EEV** | Extracellular enveloped virions | Infectious morphogenic form of poxvirus; IMV-like particle that is further wrapped and processed and actively released from the cell; can be defined by antigen-specific neutralization and more specifically by anti-Mab c6C neutralization, lack of neutralization by anti-IMV antibodies, and/or envelope protein content | (34, 36, 38) |

EV are thought to have properties that differ than the MV that allow more efficient long-range spread. These include having relative resistance to complement, immune stealth, and efficient infection and entry kinetics(40, 41). EV neutralization mechanisms involve more complex interactions of different antibodies and involvement of complement(42, 43). It also has been reported that VacV EV show reduced susceptibility to host complement that is homologous to the cell line in which the EV was propagated(44). The extent this applies to other poxviruses is still unknown, as clade I MPXV EV showed little resistance to the effects of complement (45). VacV EV are also thought to be more fragile and less likely important for inter-host transmission(46).

Depending on the strain of vaccinia virus and/or variola virus, it has been shown that at least some poxviruses either retain or release more EV virus. The increase in EV concentration typically corresponds to less retention of CEVs or the generation of released CEV, or rCEV(39, 47), Blasco, 1992 #89}. We have previously shown that we can both release and define CEV/rCEV (CEV-like) and EEV based on lack of neutralization activity or high neutralization capacity, respectively, by a specific monoclonal against EV protein A33 (39). Regardless of viral isolate, a certain percentage of CEV-like particles exist within the total composition of EV (39, 48). What makes rCEV/CEV less susceptible to neutralization by this A33-targeted antibody, the functional difference and biological role in pathogenesis (if any), stability, and/or other influences controlling their natural release have not been empirically sought. Therefore, it was one goal of the manuscript to help characterize such features as it pertains to monkeypox virus relative to the protype poxvirus, vaccinia virus.

In these studies, we utilized new and previously reported tools(39, 45) to comparatively characterize differences between rCEV and EEV, and what factors influence the composition, production and/or stability of EV that may be important for the intra- or inter-host viral cycles. We found that differences in host cells and/or isolates of MPXV and VacV influence such factors. Furthermore, we utilized oropharyngeal and blood samples that were collected from NHPs intravenously infected with MPXV clade Ia MPXV to elucidate the kinetics of EV and EV-specific antibody responses to help understand both our model, the potential function of MPXV morphogenic forms, and potential impact on transmission.

## Results

### MPXV EV Stability

#### Infectious MPXV EV is still detectable after 14 weeks under experimental conditions

The stability of MPXV EV overtime has not previously been evaluated; therefore, we asked if clade I and clade IIb MPXV EV are stable over 14 weeks at 4° C in media (**Figure 1A and 1B**). Regardless of clade, MPXV EV lost approximately 50% (mean and standard deviation of 49.2 ± 23.2% to 52.7 ± 22.2%) infectivity under MV neutralizing conditions, utilizing mAb 7D11 to neutralize anti-L1 susceptible virions (i.e., MV and membrane altered EV), after 3 weeks of refrigeration. After 14 weeks there was a decline for both clade Ia MPXV to 12.6 ± 1.34% and clade IIb MPXV to 9.13 ± 3.87%, respectfully (**Figure 1A**). However, infectious EV were still detectable at levels ranging from 500 to 9,775 PFU/mL for clade Ia MPXV EV and 50 to 300 PFU/mL for clade IIb MPXV EV at 14 weeks (**Figure 1B**). These data support that that Mab 7D11 resistant EV from either MPXV clade remains stable over an extended period under environmentally controlled conditions.

**Figure 1.**
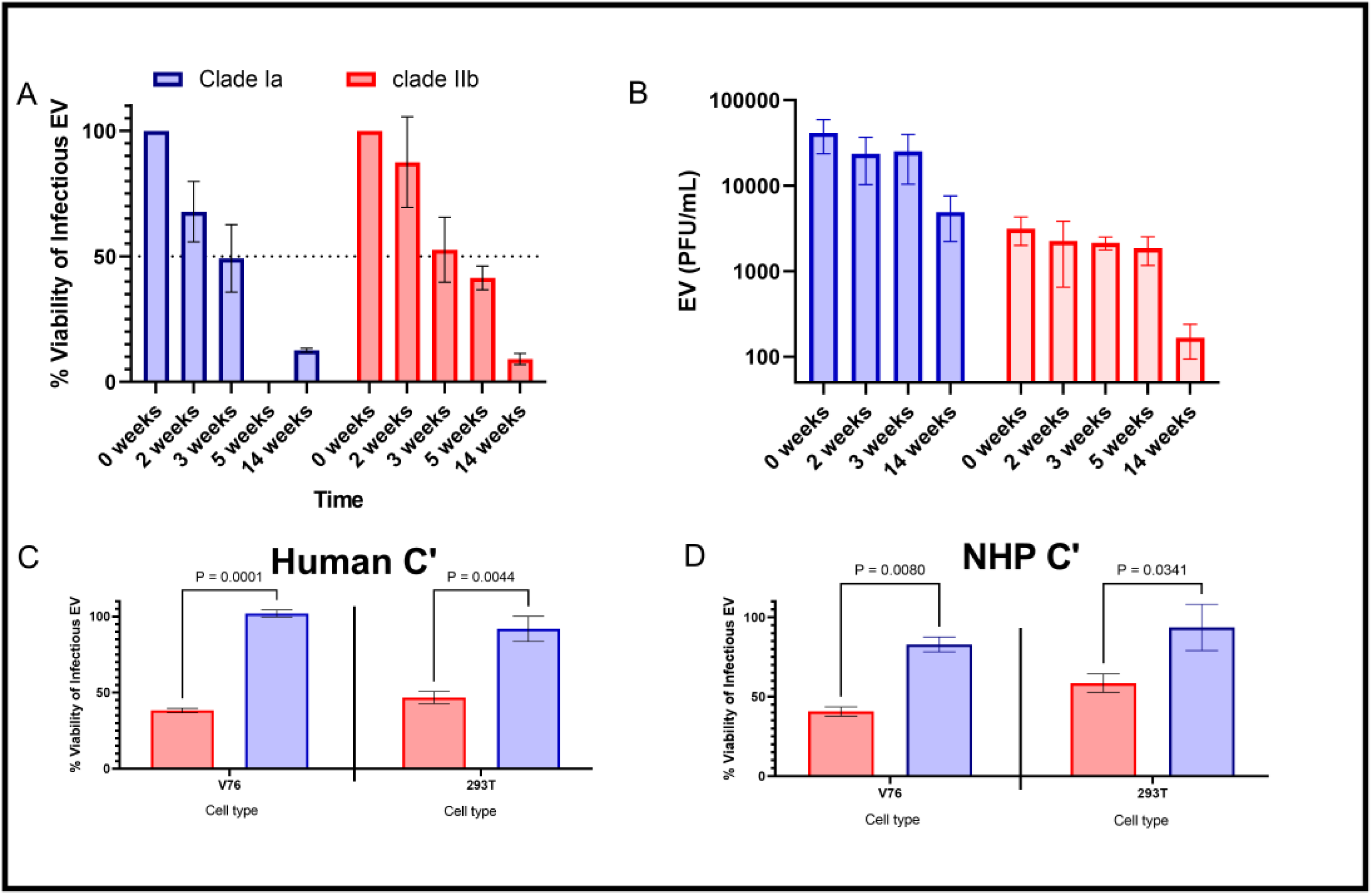
Stability of MPXV EV in two different cell lines. Preparations of both clade I and IIb MPXV supernatants from Vero 76 cells and HEK 293T cells were tested to determine both viable EV over time (A, B), as well as sensitivity to 50% human complement (C, D). 5% concentrations of the cynomolgus and human complement are typically utilized for neutralizing antibody activity and were used as a comparator (Figure 2) Initial infectious EV content of the supernatants were determined (“0 weeks”), compared to titers over time and a % viability based on titer is provided (A). Titers utilized to generate the values are also given (B). Data from both HEK293T and Vero 76 cells were combined (A, B). The % viability of infectious EV in high concentrations (50%) of human (C) and (50%) macaque (D) sera that were normalized to corresponding complement inactivated (heat inactive) human or macaque sera. P-values were calculated using a two-way ANOVA and are provided.

#### Clade IIb monkeypox virus EV are more sensitive to complement of humans or nonhuman primates independent of derivation of matching host cell type

We next asked whether MPXV EV were susceptible to host complement homologous to the cell line from which they were prepared (**Figure 1C and 1D**). EV stocks of both clades were tested for their susceptibility to 50% human or cynomolgus macaque serum or complement (heat) inactivated serum by measuring the viability measured of the remaining EV (**Figure 1C and 1D**). Regardless of cell type from which the EV was derived, the viability of clade IIb EV were compromised by the addition of either serum. Clade IIb EV were only slightly, but not significantly, more resistant when propagated from HEK293T rather than Vero 76 cells, having mean % viabilities of 46.8 vs 38.2 against human sera (**Figure 1C**) and 58.6 vs 40.7 against cynomolgus sera (**Figure 1D).** Clade I MPXV EV retained most of the initial infectivity regardless of cell lines. Vero 76 cell-derived EV had a slightly higher % viability against human serum (mean of 102 ± SD 3.90%) than cynomolgus serum (82.9 ± SD 8.08%), but opposite when EV from HEK293T cells were incubated with both serum species (91.9 ± SD 14.3 % vs 93.6 ± SD 25.5, respectively). These were not significantly different. When comparing clade I MPXV EV, viabilities were significantly higher than clade IIb independent of cell type or the two sera examined (**Figure 1C and 1D**).

To ask whether these findings were only characteristic of MPXV EV, the sera and/or cell lines chosen, we examined the complement sensitivity of VacV as well. VacV IHDJ EV were propagated from V76 and HEK 293T cells and tested against 50% (**Figure 2 A-C**) (final) or 5% (**Figure 2D**) macaque or human serum. For these experiments, we also included rabbit serum as a control, which had previously been shown to inactivate VacV EV produced in human cells in stability assays (44). Relative to HEK-293T propagated EV (HEK-EV), VacV-IHDJ EV from Vero 76 cells (V76-EV) had significantly less viability against human serum (**Figure 2A**) with similar viability obtained utilizing the VacV-WR derived from V76 cells (**Figure 2C**). The viability of V76 derived EV retained slightly higher infectivity against serum from macaques (mean of: V76-EV, 72.3% and HEK-EV, 61.0%), and lower when tested serum from rabbits (mean of: V76-EV, 23.6 and HEK-EV, 48.0%). comparison of V76 derived VacV WR EV again was similar in viability as the IHDJ strain from the same cell line. Neither of the two comparisons were statistically significant. Reducing the concentration of human serum to 5% also did not significantly change the viability of either HEK-EV or V76-EV, but reduction in concentration did significantly change neutralization by rabbit sera (**Figure 2D**).

**Figure 2:**
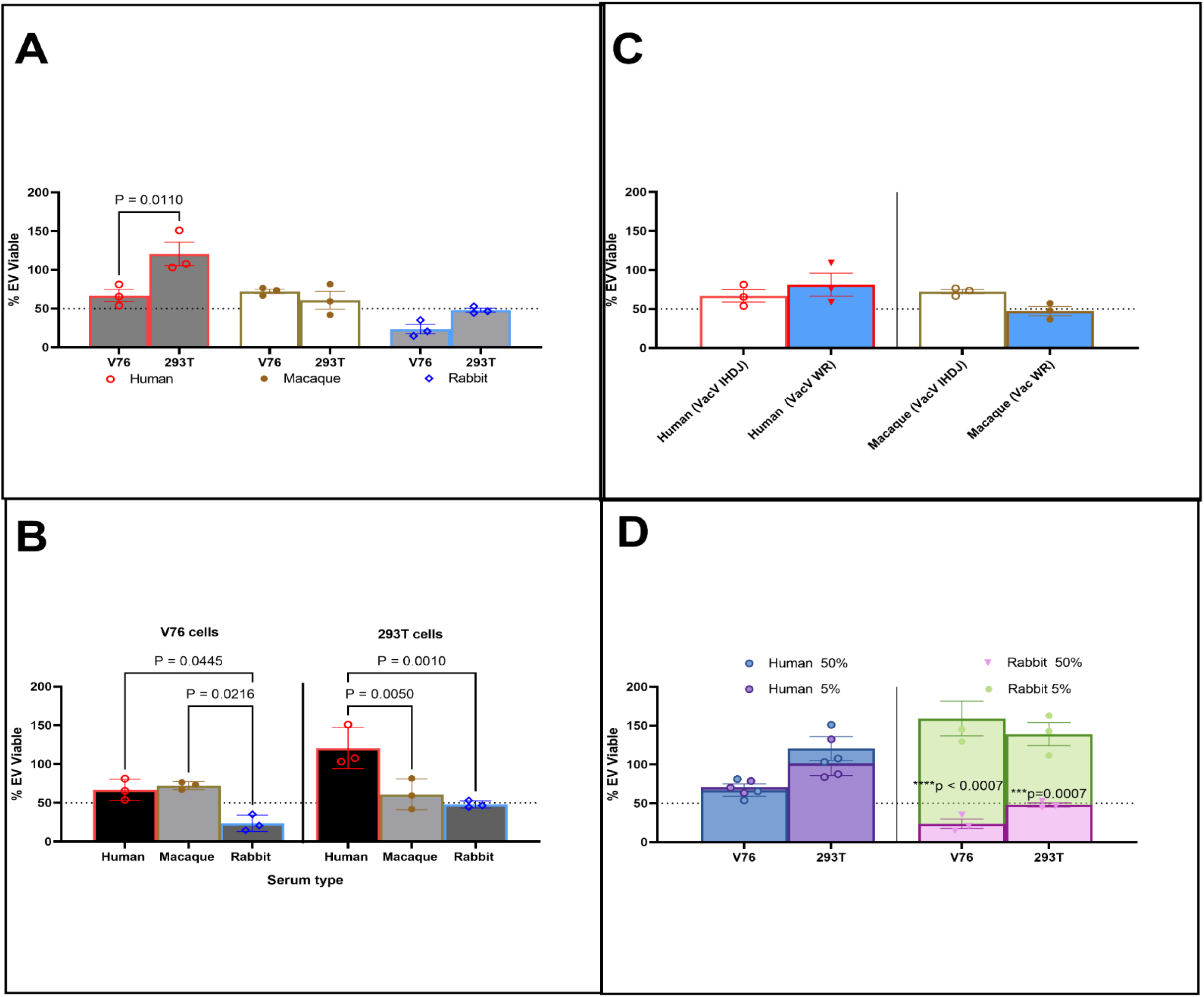
Stability of vaccinia virus EV derived from two different cell lines. Human, macaque and rabbit serum or heat inactivated serum were incubated with a target concentration of vaccinia virus EV propagated from either HEK 293T cells or Vero 76 cells: direct and statistical comparisons are shown between remaining infectious EV relative to: A and D cell type; B and C, host complement. Statistics were performed utilizing a 2-way ANOVA (Graphpad Prism).

The culmination of these studies provide evidence supporting the resistance to sera on poxvirus requires more than cellular encoded complement resistance, and likely hinges on genetic requirements, such as complement receptor mimics, of the virus. Furthermore, there is some stability of mAb 7D11 resistant MPXV and VacV EV particles, to at least 14 weeks at 4° C. (**Figure S1**) (39, 49) to A33 only)

### Characterization of MPXV EV in Cynomolgus Macaques

We next asked if and or how MPXV EV could contribute to disease pathogenesis and/or transmission utilizing samples from macaques that were exposed to intravenous MPXV clade Ia; more specifically, detect EV that were either shed via oropharyngeal swab or circulating in serum and could potentially be involved in transmission or systemic dissemination after infection and initial replication in the host, respectively.

In the first study, serum viremia was not consistent between all animals and/or days (Table 1). Concentrations of total circulating virus were low with most having typically less 100 PFU/mL or less (data not shown). Only a single sample had levels greater 100 PFU/mL (Animal 3, Day 9). Although quantifiable amounts of anti-MV resistant virus were detected in this sample, the total amount of virus was above the limited dilutions tested (**Table 1A**). No virus was detected in the sera samples from Study 2. Considering the limitations of the experiment, such as the low quantities and virus concentration of samples, the number and sampling frequency (three collection days and five total samples), the data show *trends that infectious virus protected from the neutralization, as detected by our anti-MV antibody, tended to drop over the course of disease (from 43.5% (n=2) to 0.20 % (n=1). When averaged over examined timepoints, approximately 39.5% (n=5) were not susceptible to neutralization by our anti-MV antibody*.

**Table 1:** Detection of MPXV Clade I EV in orally shed and systemic samples (sera) from experimentally inoculated cynomolgus macaques, A, and reduction of infectivity of swab samples to human complement, anti-MV Mab (7d11), or anti-EV Mab (c8A), B.

| NHP Origin | ID | 2 to 3 |  | 5 |  | 6 to 7 |  | 8 to 9 |  | 10 to 11 |  | 12 to 14 |  | 15 |  |
| --- | --- | --- | --- | --- | --- | --- | --- | --- | --- | --- | --- | --- | --- | --- | --- |
|  |  | Serum | Swab | Serum | Swab | Serum | Swab | Serum | Swab | Serum | Swab | Serum | Swab | Serum | Swab |
| Asian | 1 | ND | ND | 100% | ND | 50.00% | FA | FA | 0.31% | N/A | N/A | D | D | D | D |
|  | 2 | 12.00% | 12.00% | ND | 2.17% | ND | FA | FA | 0.18% | N/A | N/A | 0.20% | 0.20% | N/A | N/A |
|  | 3 | 75.00% | 0.00% | ND | 0.00% | 0% | FA | FA | FA | N/A | N/A | D | D | D | D |
| Mauritius | A | ND | ND | ND | 0.00% | ND | n/a | ND | 5.40% | ND | 0.51% |  | 1.08% |  |  |
|  | B | ND | ND | ND | 0.90% | ND | n/a | ND | 1.13% | ND | 0.70% |  | 1.40% |  |  |
|  | C | ND | ND | ND | N / A | ND | 0.48% | ND | 0.71% | ND | 1.55% | D | D | D | D |
|  | D | ND | 0.00% | ND | N/A | ND | 1.17% | ND | 3.19% | ND | N/A |  | 3.85% |  | 1.60% |
| AVG |  | 43.50% | 4.00% | ND | 0.77% | 25.00% | 0.83% | ND | 1.82% | ND | 0.92% | 0.20% | 1.63% | ND | 1.60% |
| MEDIAN |  | 43.50% | 0.00% | 100% | 0.45% | 25.00% | 0.83% | 1.67% | 0.92% | ND | 0.70% | 0.20% | 1.24% | ND | 1.60% |
| Overall estimated %EV-average (n) |  | **SERUM: 39.5 (5)** **SWAB: 1.7 (24)** |  |  |  |  |  |  |  |  |  |  |  |  |  |

|  | 3 |  |  | 5 |  |  | 6 |  |  | 8 |  |  | 9 |  |  | 11 |  |  | 12 |  |  | 14 |  |  | terminal |  |  |
| --- | --- | --- | --- | --- | --- | --- | --- | --- | --- | --- | --- | --- | --- | --- | --- | --- | --- | --- | --- | --- | --- | --- | --- | --- | --- | --- | --- |
|  | % reduction by: |  |  | % reduction by: |  |  | % reduction by: |  |  | % reduction by: |  |  | % reduction by: |  |  | % reduction by: |  |  | % reduction by: |  |  | % reduction by: |  |  | % reduction by: |  |  |
| Animal ID | 7D11 | C' | C' + c8A | 7D11 | C' | C' + c8A | 7D11 | C' | C' + c8A | 7D11 | C' | C' + c8A | 7D11 | C' | C' + c8A | 7D11 | C' | C' + c8A | 7D11 | C' | C' + c8A | 7D11 | C' | C' + c8A | 7D11 | C' | C' + c8A |
| A |  |  |  | 100 | 57 | 57 |  |  |  | 95 | 38 | 37 |  |  |  | 99 | -26 | 77 |  |  |  | 99 | 91 | 90 | 98 | 60 | 70 |
| B |  |  |  | 99 | 68 | 74 |  |  |  | 99 | 62 | 56 |  |  |  | 99 | 78 | 79 |  |  |  | 99 | 84 | 84 | 98 | 85 | 86 |
| C | ND | ND | ND |  |  |  | 100 | 34 | 61 |  |  |  | 99 | 64 | 31 |  |  |  |  |  |  |  |  |  |  |  |  |
| D | 100 | -16 | 5 |  |  |  | 99 | 69 | 67 |  |  |  | 97 | 88 | 89 |  |  |  | 96 | 43 | 66 |  |  |  |  |  |  |
| Average | 100 | -16 | 5 | 100 | 63 | 65 | 99 | 52 | 64 | 97 | 50 | 46 | 98 | 76 | 60 | 99 | 26 | 78 | 96 | 43 | 66 | 99 | 87 | 87 | 98 | 72 | 78 |

C.
|  | Day 3 |  | Day 5 |  | Day 6 |  | Day 8 |  | Day 9 |  | Day 11 |  | Day 12 |  | Day 14 |  | Terminal |  |
| --- | --- | --- | --- | --- | --- | --- | --- | --- | --- | --- | --- | --- | --- | --- | --- | --- | --- | --- |
|  | 7d11 minus C' | (C'+c8A) minus C' | 7d11 minus C' | (C'+c8A) minus C' | 7d11 minus C' | (C'+c8A) minus C' | 7d11 minus C' | (C'+c8A) minus C' | 7d11 minus C' | (C'+c8A) minus C' | 7d11 minus C' | (C'+c8A) minus C' | 7d11 minus C' | (C'+c8A) minus C' | 7d11 minus C' | (C'+c8A) minus C' | 7d11 minus C' | (C'+c8A) minus C' |
| Difference in Neutralization and C' addition (%) | -116 | 21 | -37 | 2 | -47 | 12 | -47 | -4 | -22 | -16 | -73 | 52 | -53 | 23 | -53 | 0 | -12 | 6 |
| % EV Estimates | 1.6** |  |  |  |  |  |  |  | 10.7 * |  |  |  |  |  |  |  |  |  |

\*Data utilized to calculate: Difference in Neutralization (average of red values from Table B)
\*\*Data from Average 7d11 (blue blocks from average Table B) and subtracted from 100%.

Infectious virus from oral swabs were run “fresh” in the absence of sonication and/or freeze thaws and within approximately one week of collection. For both Study 1 and 2, infectious virus was detected more consistently between animals, and much more abundant per sample than sera. The detection for individual swab samples resistant to anti-MV antibody varied between 0 and 12% (**Table 1A**). The median value tended to be closer to <1%, with a general increase from 1.24 to 1.60%, increasing relatively to the day collected post exposure (**Table 1A**). These data suggest that EV-like particles that are resistant to anti-MV antibody are present in oral shedding(s) in quite small percentages and tend to be higher very early in disease, after the onset of virus shedding (typically day 3+).

For the second study, concomitant with the above assay design, we also asked whether the infectious material from the oral swabs was sensitive to the effects of human serum or if we could assess EV titers in a slightly different manner, e.g., an anti-EV mAb under neutralizing conditions with mAb c7D11 and complement (**Table 1B**). Because the anti-EV antibody requires complement to neutralize EV efficiently (39) and both complement and or anti-MV antibody (mAb c8A) can neutralize a proportion of released EV from MPXV (45), it was necessary to analyze the individual components involved to discern potential intact EV from MV or MV-like infectious particles. Anti-MV neutralization was also included, extrapolated from **Table 1A**, as a comparator for total intact EV.

Other than Day 3 post exposure, swab supernatants were sensitive to the effect of additional human complement. From all samples tested, addition of human complement increased titers of two of the 16 samples tested relatively 16% and 26% suggesting no neutralization; our reported increases in neutralization of the remaining 14 samples ranged from 31-91% (average of 65.8%, STD of 18.5% and median of 66%). No obvious trends based on time/day of collection were noted but did range from no neutralization on Day 3 (n=1) to 87% on Day 14, with many fluctuations in between.

When the EV neutralizing antibody was also included with the complement, most differences between complement alone were small with the exceptions of Day 11 sample from Animal A and Day 3 sample from Animal D, for which no neutralization by complement was detected (**Table 1B**). Anti-B5 (complement + mAb c8A) calculation of EV (when complement neutralization is subtracted out) ranged from an overall % average of EV that was less than 0 (e.g., no EV detected) on Days 8 and 14 to a high of 52% on Day 11, again with no discernable trend relative to Day of infection (**Table 1C**,). The overall average of %EV was calculated to be 10.7% average of all days analyzed. These data support the low MV-based neutralization resistance percentages (to mAb 7D11), and that there are relatively few complement resistant EV-like particles neutralizable by anti-B5 (mAb c8A) antibody relative to those that have neutralizing properties of MV-like particles (mAb 7D11).

Since little to no infectious virus detected in our assays for study 2 using cell free serum samples, we questioned whether there was any evidence that the virus might be present, but non-infectious. To test, we conducted qPCR testing on the serum (free virus) samples of animals A, B, C and D from study 2. Whole blood was also analyzed by PCR to represent total potential virus (cell associated virus) and compared to the genome copies estimated virus concentration (ePFU/ml) in the serum (**Figure 3**). Virus genetic materials were first detected either concomitant (Animals A-C) or earlier (Animal D) in serum relative to whole blood. Although the magnitude of detected genomes was initially similar (>6 days), whole blood had overall greater quantities of detected nucleic acid.

**Figure 3.**
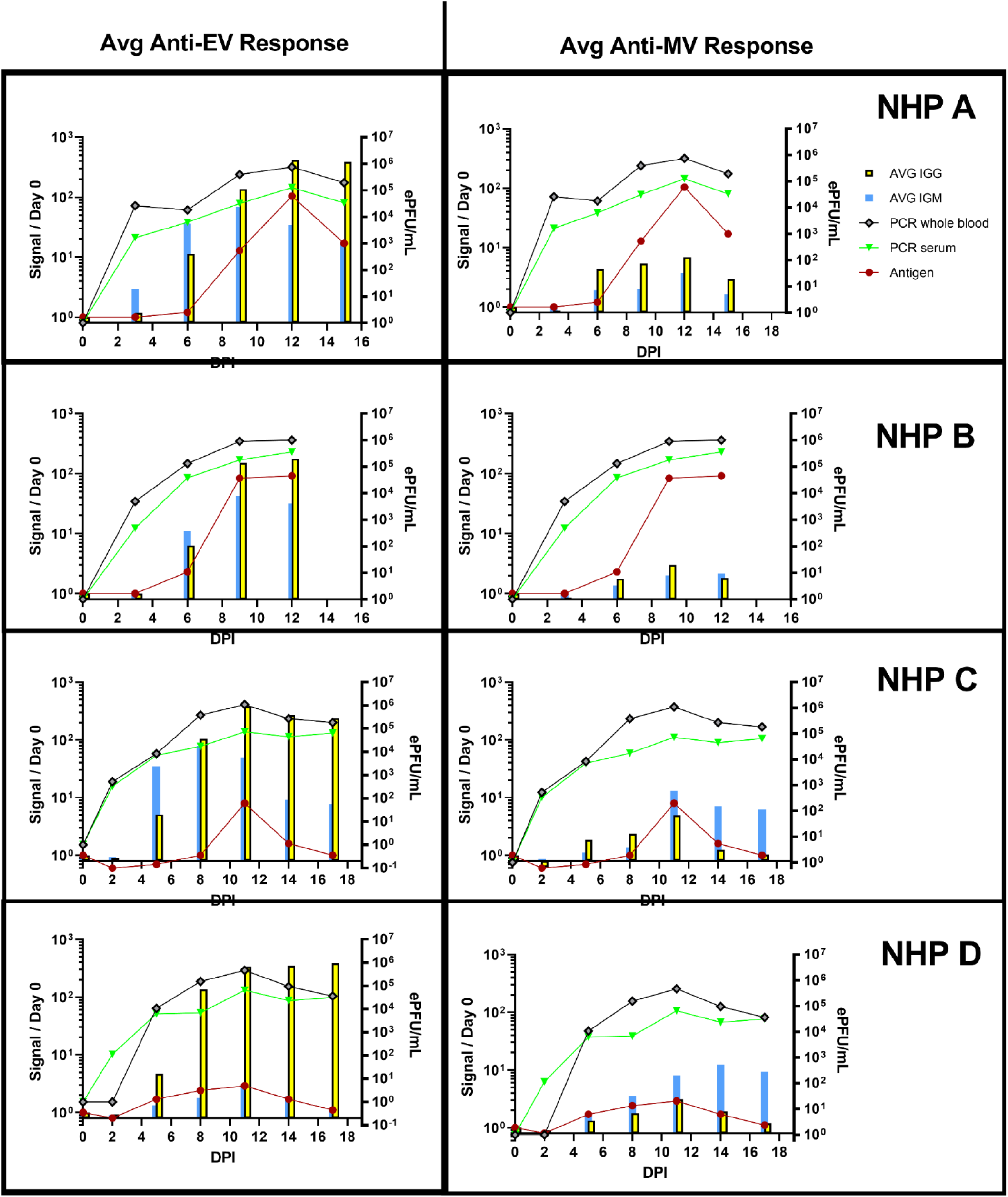
Characterization MPXV detection and morphogenic specific antibody response samples from NHPs exposed to intravenous clade I MPXV. Whole blood (grey lines) and serum (green lines) from animals sampled (**see Table 1**) were analyzed by PCR and normalized into an estimated PFU (ePFU per mL) based on quantity of genome (right Y-axis in all graphs). MPXV antigen detection (red lines), as well as host specific IgG (yellow bars) and IgM (blue bars) to VacV and MPXV EV (left column) and MV (right column) antigens, were also normalized to signal from samples acquired prior to exposure on Day 0 (Signal/ Day 0; left axis of all graphs).

Since serum contained detectable nucleic acid sequences of MPXV, we asked whether we could detect more evidence of particles circulated systemically. To do so, we tested the serum samples for the presence of poxvirus protein antigen by Magpix (**Figure 3**). Detection of antigen was noted in all animals by Day 6 post-MPXV exposure. Antigen increased the entire disease course in 2 of 4 animals (**NHP A and B**) and all had peak antigen titers between Days 10 and 12 post exposure (**Figure 3**). Animals C and D declined after peaking. The culmination of nucleic acid and antigen in the serum suggest that viral particles may have been present but were not infectious or infectious at undetectable quantities.

We have already explored MPXV-EV neutralizing effect of antibody specific for MV (45), EV(50), and/or complement(45), as well as reported in this manuscript. Next, we investigated the contribution of other factors, such as MV and/or EV specific antigen specific antibody responses and determine how they may play a role in the systemic viral kinetics of the intravenous-NHP model (**Figure 3**). In the same serum samples as MPXV antigen and genome detection, we looked for antibodies specific for EV antigens against MPXV (A35 and B6) and VacV (A33, B5); as well as MV antigens MPXV (A29) and VacV (A27 and L1). IgM antibody to EV was detected by Day 3 in 2 of 4 animals (**Figure 3**). Specifically, NHP D that was sampled on Day 2 and NHP A on Day 3 had a two- or three-fold average increase than overall background, respectively. Similar or greater changes of IgM against the VacV and MPXV MV antigens were not detected until Day 5 or 6 post intravenous exposure. Detection of IgG against EV antigens occurred in all animals on the same days with a range of 4.7-10-fold relative increase in detection (**Figure 3**). In all four NHPs, IgG and IgM against EV targets hit peak values on sampling Days 11 and 12 and IgG levels maintained constant for the remainder of the study in life with IgM slowly declining in magnitude. Antibody levels of IgM and IgG against MV targets were slightly different in kinetics than EV targets, peaking on Day 9-12 and IgG then declining in all animals (**Figure 3).** Interestingly, IgM against MV antigens remained relatively unchanged or increased in 3 of 4 animals (NHPS B-D) until the end of the study.

### Virus and cell specific EV changes: EEV, CEV, rCEV

#### rCEV and mAb c6C-induced rCEV are capable of being neutralized by a different A33 antibody

It has been shown that mABs against the target A33 may not be fully effective for neutralizing all potential poxvirus agents of interest either because of lack of binding(49), binding to A33 but only neutralization specific morphogenic forms (i.e., mAb c6C {Mucker, 2020 #1071), or if A33 was/is not present on the EV particle. If so, a combination of either A33 directed or cocktail against different antigens is necessary for broad protection and decreasing the resistance to the therap(ies).To answer whether another anti-A33 targeting mAb could supplement the activity of mAb c6C and determine if the A33 protein is abrogated after mAb binding and/or missing from CEV particles, we first had to demonstrate if a monoclonal antibody could neutralize both EEV- like and CEV-like forms. Therefore, we first tested two previously reported antibodies, mAbs 10F10 (**Figure S2**) and 1G10 (Figure 1A), against both VacV International Health Department, isolate-J strain (IHDJ) and VacV Western Reserve strain (WR) for the capacity to neutralize EV in the presence of human complement. Consistent with previous reports mAb c6C was only able to neutralize 50% (average of 50.0 -54.2%) of the VACV IHDJ EV at the highest concentrations, but 90-95% of VacV WR EV. MAb 1G10 was able to neutralize 95.4 -97.0% of the same stocks of virus. These data suggest that mAb 1G10 may neutralize the entire EV population of the supernatants. 10F10 had a neutralizing profile similar-to mAb c6c, as it neutralized up to 99% of the EV from VacV WR but had limited neutralization capacity (≤ 59%) against VacV IHDJ (**Figure S2**). For these and other reasons (see discussion), mAb 1G10 was utilized for further experimentation.

Because rCEV-like particles are resistance to neutralization by mAb c6C, next we asked if mAb c6C effectively blocked or disrupted A33 present on rCEV-like particles as to hinder A33- directed mAb 1G10 neutralization of the particles. To differentiate rCEV-like particles from EEV stocks were defined by susceptibility to neutralization by mAb c6C or an anti-B5 antibody (mAb c8A), respectively. VacV WR virus was propagated in the presence of a dilution series of mAb c6C for approximately 2 hours, then either mAb1G10, additional mAb c6C, or media alone was added to assess virus neutralization of the release virus (**Figure 4B**). There was an inverse relationship of concentrations of mAb c6C in the inoculum to the titer of rCEV-like particles produced in vitro (**Figure S1**). In this experiment, the effect of additional mAb c6C was used to as a control for any further total CEV release or mAb binding/neutralizing of the total EV (**Figures 4B and Figure S2)**. The addition of mAb c6C to the rCEV had little to no change in the neutralizing effect at dilutions at or less than 1:250 (0.243 ug/mL) but trended higher relative to reduced concentration (**Figure 4B**). In contrast, regardless of mAb c6C concentration used in the preparation of the EV, the addition of Mab 1G10 to the inoculums caused significant changes in neutralization relative to Mab c6C with greater than 50% of the Mab c6C treated EV virus preparations being neutralized with mAb 1G10 (**Figure 4B**). These data suggest that A33 is functionally present on virions rCEV after treatment with mAb c6C and the mechanism required for A33-based neutralization by antibodies is not hindered, regardless of whether the rCEV-like particles are artificially induced by mAb c6C or inherently produced by VACV-IHDJ (**Figure S2**. Therefore, appropriate supplement of another mAbs targeting A33 could potentially help increase the protection against a breadth of strain variations.

**Figure 4:**
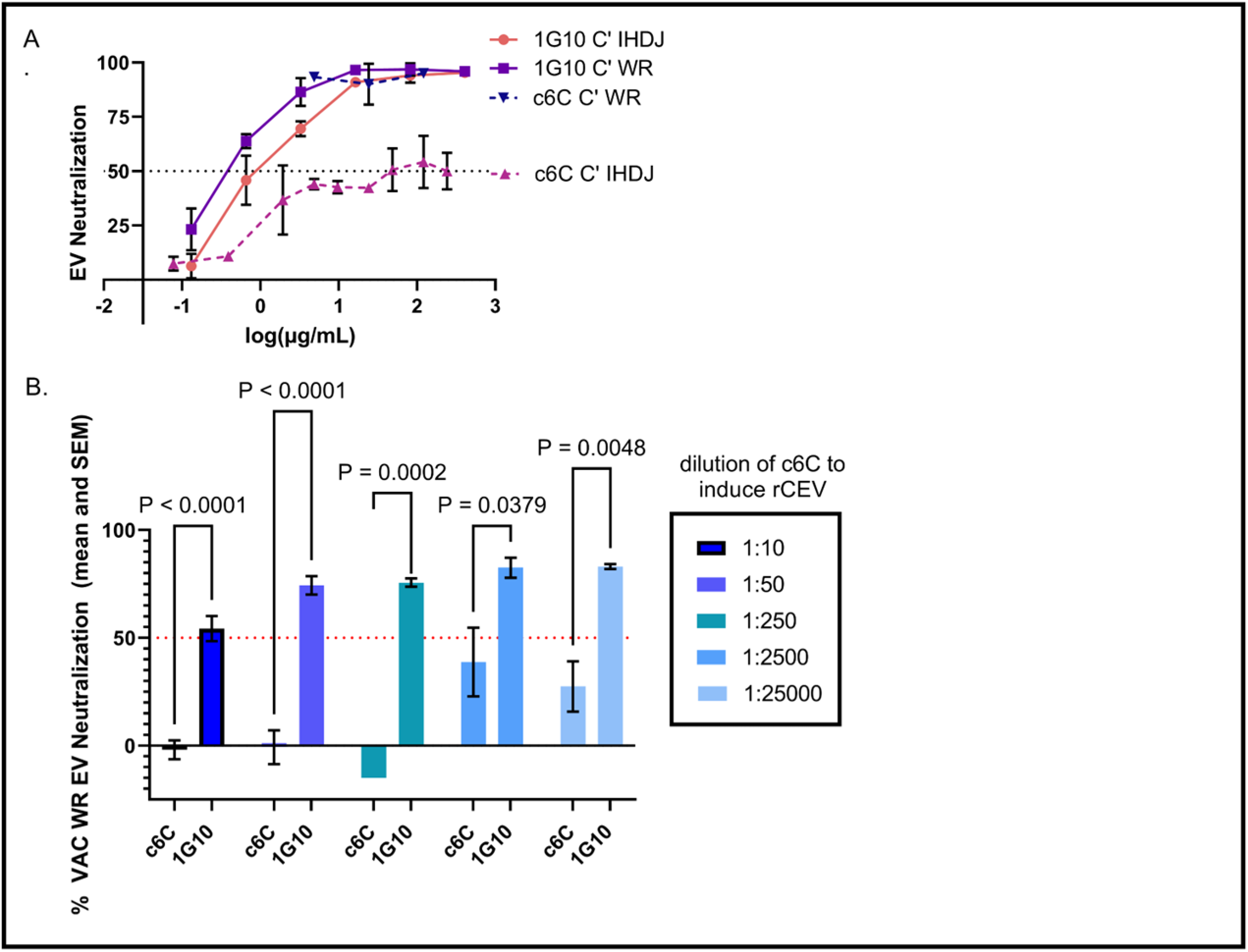
Neutralization of vaccinia virus EV using ant-A33 antibodies of different mechanisms. EV from strains WR (A, B) and IHDJ (A) of vaccinia virus were assayed for infectivity in the presence of Mab 1G10 or anti-EEV antibody c6C in the presence of complement and anti-MV antibody 7D11 (A). CEV were released (rCEV) from cells utilizing different dilutions of c6C (see legend), a constant concentration of Mab 1G10 was added to each and EV neutralization were assessed. A two-way ANOVA with Tukey’s multiple comparison was utilized, and significant P values are given.

#### Cell line can affect VacV EV output and composition

Since similar strains of vaccinia virus generated distinct EV populations, it was possible that cell type(s) (i.e., related to specific host-virus cell interaction) would influence the composition of the extracellular population of virions. We have previously utilized Vero cells to produce and study clade I MPXV EVs, and anecdotally noted an increase in output relative to VACV-WR EV produced in HEK 293T cells(39, 45). Therefore, we questioned whether infection of Vero 76 cells would increase our output of VacV WR EV and/or alter the composition of the EV produced. Cell lines were infected with either the WR or IHDJ strain of vaccinia virus. For the WR strain, extracellular virus statistically increased (p=0.0007) by two logs when produced from Vero 76 cells relative to HEK 293T cells under the same conditions **(Figure 5A).** When grown in HEK 293T cells the IHDJ strain of VacV had statistically higher EV titers than those produced from either cell type of WR (p < 0.0001 HEK 293T EV and V76 p=0.0186); whereas there were no differences between virus strains when both were grown in Vero 76 cells. These data suggest a difference in cell type and strain could affect production of EV when using similar multiplicity of infections and consistent inoculums.

**Figure 5.**
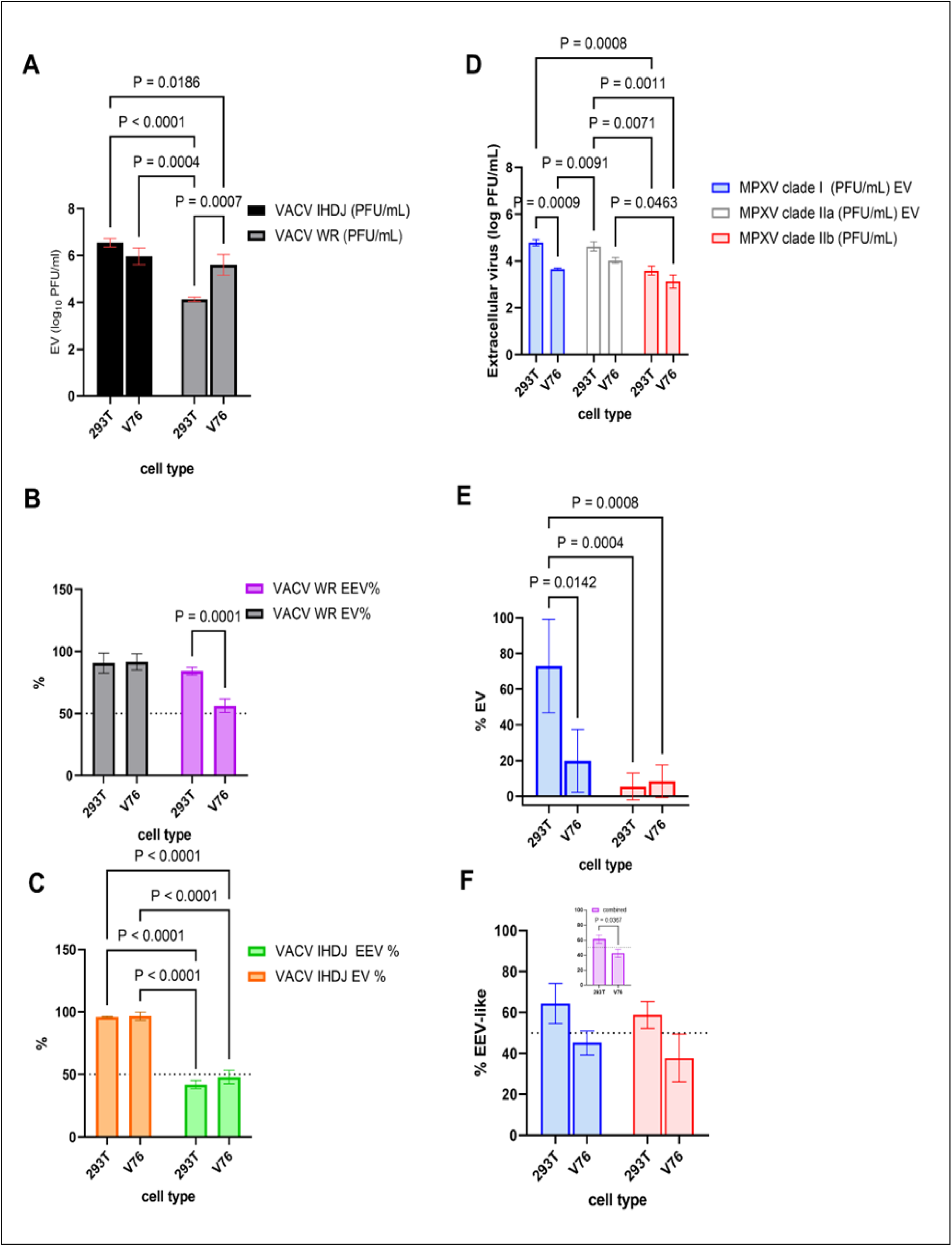
Differences in EV composition between two cell lines using strains of MPXV and VACV orthopoxviruses. Vaccinia virus strains WR (A, B) and IHDJ (C), and MPXV clade I and clade II (D, E, F) representatives were grown on either HEK 293T cells (293T) or Vero 76 cells (V76), supernatants titrated in the presence of MV neutralizing antibody, and titers (A, D) were used to calculate EV content (%EV) based on percentage not neutralized by anti-MV antibody (B) and confirmed by antibody neutralization by anti-B5 antibody (B, C) or to calculate percentage EEV-like particles (%EEV) based on resistance to neutralization of Mab c6C (B, C, F). Data from MPXV combination of both clade Ia and clade IIb is shown in the inset of F. Mean and standard deviations are shown.

We next wanted to both confirm the EV-like nature of the released virus and, more specifically, whether the functional (infective) proportion of CEV-like and EEV-like virions varied between different cell lines and/or vaccinia strains. Harvested supernatants were subjected to neutralization by an anti-MV antibody and human complement to determine the percent of the total supernatant population of EV and confirmed in the presence of anti-B5 antibody, now termed % EV (**Figure 5B and 5C**). The same supernatants were also subjected to mAb c6C, to define differences between EEV-like and CEV-like EV for both strains of VacV. Although there were no significant differences between cell lines based on the proportion of VacV WR EV susceptible to anti-B5, another EV specific protein, neutralization, the number of mAb c6C susceptible EV produced were significantly fewer when produced in Vero 76 cell line relative to HEK 293T cells (**Figure 5B**). When repeated using the IHDJ strain of VacV (**Figure 2C**), there were significantly fewer Mab c6C susceptible particles (EEV) predicted by B5 neutralization regardless of cell line. Therefore, EEV composition can be altered given the infection of different cell lines for WR strain, but not the IHDJ strain of VacV.

We then used serial timepoint harvesting of both cell types infected concomitantly with VacV WR and evaluated by transmission electron micrography (TEM). Vero 76 cells showed formation of relatively more virions around the outer plasma membrane compared to HEK293T cells at the same time points, consistent with more CEV production in these cells (**Figure 6A**). Also, a lack of comet phenotypes in HEK293T cells, but a large increase in relative plaque size and spread (i.e. EV production) in Vero76 for both VacV IHDJ and WR, as can be observed by comet assays (**Figure S2)**. Together, both the micro and macro-imaging support the findings by the neutralization-based assays that the two cell lines are producing different proportions of EV-like particles. Therefore, the combination of data from these experiments shows that for VacV strain WR, changes in cell type alone can alter the quantity of EV-like progeny that contain significantly different compositions of rCEV-like and EEV-like subtypes.

**Figure 6.**
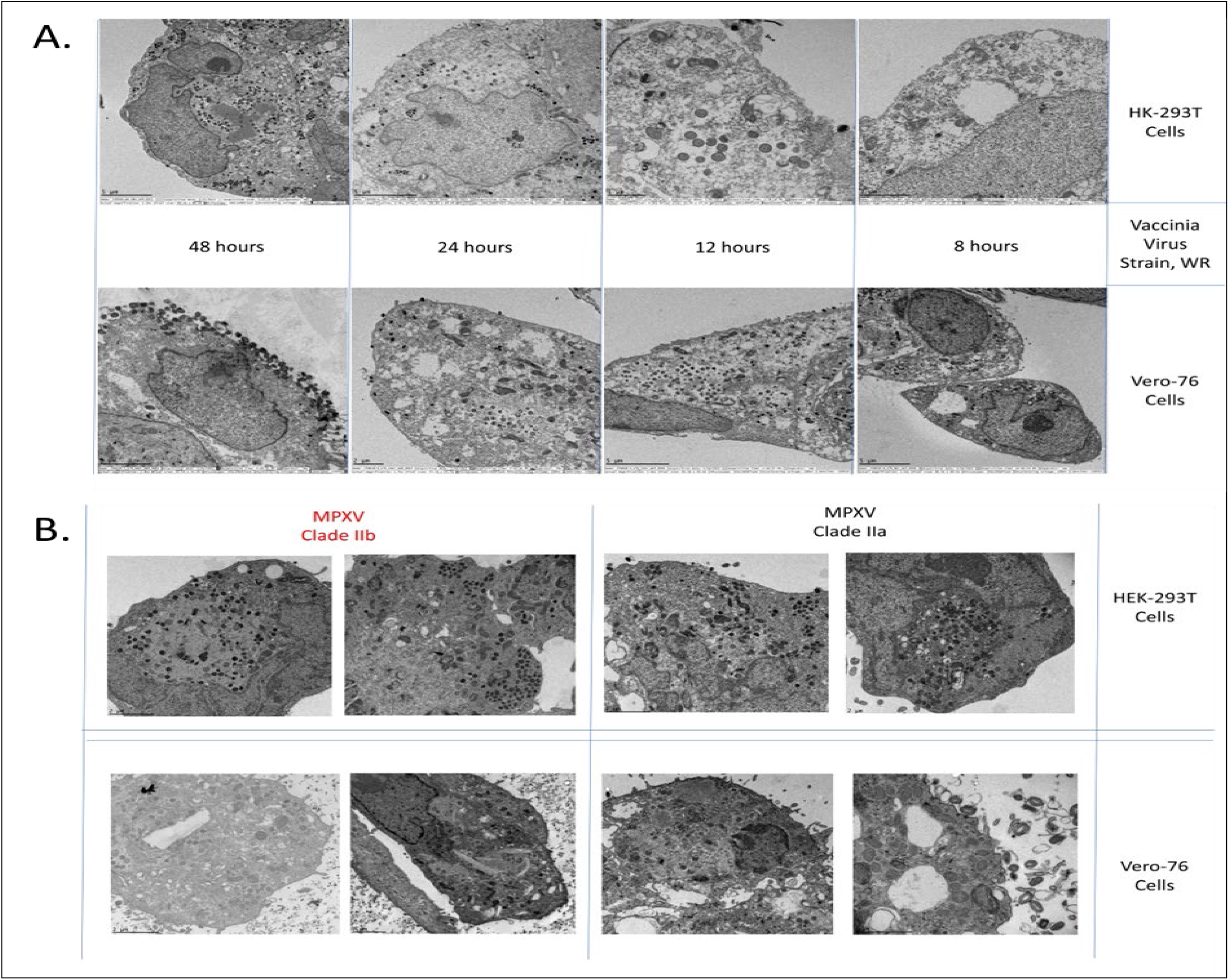
Transmission electron micrographs HEK-293T and Vero 76 cells. HEK-293T or Vero 76, top and bottom panels of A and B, respectively, were infected with the same preparation of VacV strain WR (A) or two different clades of MPXV (B) and inactivated. Representative images of VacV are shown at the various time points in hours post infection, whereas MPXV is shown at 48 hours (B).

#### Cell line selection can also affect MPXV EV output and composition

We hypothesized that MPXV and/or specific clade I and II isolates would produce different proportion of EV in a cell type dependent manner consistent with VacV WR. Therefore, we assessed supernatants containing EV from MPXV representing clade Ia, IIa and IIb propagated in both HEK293T or Vero 76 cell lines for total EV, an estimation of % EV utilizing an anti-B5 antibody, and mAb c6Ct o estimate the susceptible or resistant %EV (**Figure 5 D, E, F).**

In terms of infectious extracellular virus, HEK293T cell line tended to produce more infectious extracellular virions for all three MPXV virus strains (**Figure 5D**). Statistically, this trend was only significant for the clade I virus, Zaire (p=0.0009). Between virus clade/subclade, extracellular virus of MPXV clade Ia produced in HEK293T cells tended to be the highest with a mean of 4.8 log(STDEV of 0.233) and significantly more than MPXV clade IIb produced in HEK 293T. MPXV clade IIb produced similar amounts of infectious virus regardless of cell type. MPXV virus clade IIa tended to be more like clade Ia under all variables, with the exception that, unlike clade Ia, there were significant difference in titer between clade IIa and IIb when grown in Vero 76 cells.

We next examine the effect of cell type and/or clade on the composition of infectious supernatants, as it pertains to resistance or susceptibility anti-L1 antibody (e.g., intact EEV or rCEV respectively). We hypothesized that analogous to experiments with VacV, that the quantity (**Figure 5D**) and/or composition (%) (i.e., EEV and rCEV-like) would differ between cell types. Therefore, we used our approximation of % composition and asked what percentage of the total infectious virus were resistant to neutralization against an anti-MV antibody (**Figure 5E**); while also testing the susceptibility of the EV-like population utilizing Mab c6c (**Figure 5F**) to further characterize the composition. We focused on clade Ia and clade IIb for this further characterization, as clade IIa was like that of clade Ia. Isolates from IIa are also a safer and practical option than the more virulent Ia isolates.

Between the two clades/subclades examined further, clade IIb produced the least amount of total anti-MV resistant EV in both cell types (**Figure 5D**). For both clades, Vero 76 cells tended to produce proportionately less anti-MV resistant particles, but only clade Ia significantly produced titers that were different to HEK293T cells within (P=0.009). When compared between clades, clade Ia and IIa produced in HEK 293T cells had significantly higher titers relative to clade IIb titers generated using either cell type (**Figure 5D**). Although higher EV titers were observed for clade Ia and IIa than IIb grown in Vero 76 cells, differences were only significant between titers of IIa and IIb (P=0.0463).

The total EV proportion (%) was also determined by mAb specific, plaque neutralization (**Figure 5E**). Clade Ia MPXV had significantly lower proportion of the total EV content when propagated in Vero 76 cells relative to similar preparations in HEK293T cells. Comparison between the two clades showed the overall percent of intact EEV was significantly less in clade IIb relative to Clade Ia in both Vero 76 (8.43% vs 19.9%, respectively) and HEK293T (5.46% vs 72.9%, respectively) and when cell lines were cross compared (**Figure 5E**).

Assays were conducted to further delineate the composition of infectious EV populations into EEV-like and rCEV-like virus; that is, mAb c6C neutralizable vs. resistant particles, respectively (**Figure 5F**). For both clade Ia and IIb MPXV, the most neutralization was detected against HEK 293T propagated viruses having similar amounts of neutralization (mean of 64.3% vs 58.9%). Again, preparations produced in HEK 293T using either clade resistance to Mab c6C had a similar neutralization (mean of 45.2% vs 37.8%). Neither differences in clade nor cell type were significantly different. When data were combined, the overall trend between cell types was significant for MPXV EV (inset of **Figure 5F**).

Cells were also infected with either clade IIa or IIb and evaluated by transmission electron micrography. Since it had a similar profile as Ia (**Figure 5D** and **supplemental Figure 3S**), clade IIa was used as a comparator in these experiments instead of clade Ia. Like VacV, observations of MPXV particles in Vero 76 cells show more total MPXV on the external cell membrane, and virus more consistent with CEV-like particles. This is far less evident in clade IIb infected cells (**Figure 6B)**. Regardless of clade, more intracellular virions were observable clustered in the cell periphery of HEK 293T cells relative to Vero 76 cells.

## Discussion/Conclusion

In these experiments, we functionally assessed the extracellular enveloped virions as a non-homogenous entity. Not only do the findings contribute to our basic understanding of poxvirus virology of poxviruses of concern, but they also strengthen how we assess countermeasures against such viruses. We found that composition, both production and characteristics, of EV-like virions are influenced by host cell line and/or specific strains of MPXV and VacV examined. Characteristics examined included stability over time and sensitivities to complement. We identified reasons for caution when developing prophylactic or therapeutic mAbs to poxvirus, primarily from the perspective of A33 targeting antibodies. We also provided granularity concerning the nature of EV and related host response(s) in a pinnacle model of mpox and smallpox disease(s).

The prevailing theory is that the overall perceived stability of poxvirus EV the particle less suitable due to inherent fragility of the outer membrane. The eventual breach of the outer envelope becomes evident under purifying condition (51)Brennan, 2020 #1162} and, in our studies, large increase in susceptibility to anti-MV antibody in a relatively short period of time (**Figure 1 A, B and Figure S1**). Given the stability data of the MPXV EV over the longer range and our finding that at least some EV-like remain productively infectious (i.e., resistance to anti-MV Mab) suggest that they should not be overlooked for their importance in transmission, but mores specific advantages of the EV relative to MV should be explored. Other factors, such as the type of particle that is mainly being by the host, as well as the sample type and timing relative to disease state should also be considered. Understanding these parameters require empirical modeling, but also requires characterization of disease in the host of interest. Others have shown MV-like MPXV clade II to be stable under environmental conditions(52).

We next questioned how the *in vitro* stability data, or relative lack thereof, empirically fit with disease. More specifically, we questioned the role of MPXV EV in viremic spread given the complement sensitivities and transmission given both complement sensitivity and temporal robustness in the intravenous macaque model of MPOX and smallpox diseases. The role of EV in systemic spread has always been questionable and is likely to be dependent on the relationship evolved between a specific virus with a specific host (45, 51); therefore, we also examined humoral host responses to EV and MV (**Table 1 and Figure 3**). We show that levels of detected viral genome continued to increase even when antigen was decreasing. This suggested that either noninfectious viral material was being released into the sera and/or infectious viral particles were being rendered inactive by innate and/or adaptive response mechanisms. We have previously shown that large proportion of freshly released EEV is susceptible to either anti-MV neutralizing antibody by neutralization assays, anti-MV antibodies and/or complement by comet reduction assays, (45). In this study, increasing complement to 50% was less impactful against MPXV Zaire relative to clade IIb MPXV EV-like infectious particles. It should be stressed that input MPXV EV titers were first defined initially in the presence of both 7D11 and 5% human complement, thereby the dose is based on selecting out at least some susceptible EV-like particles (**Figure 1 C and 1D**). One might think that most, if not all, infectious particles may be selected out and the remaining virus should be impervious to the effect of additional complement, but experiments with vaccinia virus suggest that increasing to such a level could still have a statistically significant impact (**Figure 2 D**). This is also realized when noting that with little exception (see **Figure 1 B, HEK 293T derived EV and human complement**) additional complement does impact the infectivity of the stock EV in almost all cases on a relative scale (**Figure 1 and 2**). Whether cells infected by virus derived from macaques produce adequate quantities of complement resistant EV to have any impact on disease or transmission is unknown but remains unlikely. It is also possible that macaques are inherently and evolutionary responsive to MPXV, as shown by the rapid IgM and IgG response to select EV and MV antigens (**Figure 3 left and right panels**). The onset of EV IgG response was >5 days in all animals, whereas the MV response required 5-6 days to detect. EV titers also trended towards the typical class switching from IgM to IgG over time (**Figure 3 left panel**), whereas this does not occur for MV responses in 3 of the 4 animals (**Figure 3 Right panel**), suggesting a more rapid, vigorous, and complete humoral response to EV in NHPs. Artificial differences inherent to the assay (different proteins and folding, antibodies, reagents, etc.…) may explain this as well as magnitude differences, but assays utilizing at least two MV and EV target proteins as well as vaccinia virus homologs were included to add robustness, increase sensitivity, and increase confidence in detection. The culmination of these data suggest a lack of importance of the EV in spread (and possibly disease in general) in the macaque model of smallpox and MPOX; but what we have learned throughout the experiments contained within this manuscript is that there should be some caution broadly ascribing a model of EV function until specifically confirmed under variable conditions (e.g. infection routes, strains/orthopoxviruses) in a given macaque species, let alone applying to other hosts or other hosts with similar variables.

We found little evidence that MPXV EV-like virions and/or non-cell associated virions in general were likely important for additional systemic/viremic dissemination in the intravenous NHP model system. These animals tended to have varying aspects of hemorrhagic disease equating to a more severe disease course (data separate and outside the scope of this manuscript as presented)(53). It is quite possible when infection outpaces host immune control that virus can more readily be found circulating in these animals. One can envision such a scenario when the virus is highly adapted to the host and specific immune defenses, such as smallpox in humans. Although we showed that clade Ia MPXV is more resilient to the effects of complement than our IIb isolate, suggesting some the possibility for modest stability in the blood environment (**Figure 4B**), there are some caveats. More specifically, it was previously shown that there were still a large proportion of total EV that were neutralized by either anti-MV Mab or complement(45). Again, it should be noted that we had already pre-selected EV that were resilient to use in the neutralization assays. The susceptibility of MPXV clade Ia to complement and anti-MV was also recapitulated using in vivo sampling of oral swabs and comparing two methods for calculation of EV content (**Table B**). The addition of consistent anti-MV antibody reduces the total infectious content of EV from an average of 10.7% down to 1.6% (**Table B**), establishing an EV range falls within previously reported data(45). Here data suggest that oral shedding is overwhelmingly MV in quantity and likely a large component of what would be shed in this model system. Head-to-head comparisons of MV, EV and the underlying MV-like particles (37, 54) for infectious potency or environmental stability but testing all particles were beyond the scope of our current studies. There is a requirement for such studies before totally discounting the potential contribution of EV in transmission.

Although useful for future experimentation and generating deeper questions, assays such as ours are quite rudimentary and require refinement for in vivo applications. For instance, it is difficult to ensure that we are not 1). detecting residual inoculum early in disease 2). exacerbating potentially false positive or negative results due to low titers, limited sample, and/or paucity of the sample 3). Sample integrity and collection methods can be an issue when assessing a more fragile envelope. Although functional and very biologically relevant, our assay is heavily dependent on the integrity of samples and preservation state of the infectious particles, and overall is a somewhat indirect assessment. More specifically, we get one chance to characterize in vitro samples with a two-week timeframe. After, we must consider that a portion of infectious viral population of the sample is at least somewhat altered relative to its initial collection (**based on Figure 1**). Understanding these and other limitations of our assay and applying them to our intravenous model, our findings suggest that it is likely that the requirement for high intravenous doses of any form of infectious MPXV are required for systemic (severe) disease because of the sensitivity and instability of the virus to nonspecific factors of the host environment, such as complement. *Therefore, primarily localized replication occurs and a robust secondary viremia cannot naturally be established when the virus is administered by other routes or by lowering the intravenous dose of MPXV virus. This can also be visualized by the sharp drop in lesion severity (and overall disease severity) associated with a log or more reduction in virus* (*45, 55*). Again, this result should only be considered as defined by the conditions of our studies and should not be broadened to all disease models or all poxviruses.

Infectious rCEV maintain A33 active sites and/or specific epitopes needed for entry and neutralization (**Figure 4**) by implementing A33 mAbs that performed in slightly opposing ways in screening assays. During the same screens we also discovered that mAb 10F10 had neutralization and plaque phenotype profiles like those of mAb c6C; therefore, it was likely not a good candidate for such questions. Antibody 1g10 showed the capacity to neutralize all VacV EV-like virions of both IHDJ and WR strains **(Figures 4 and S2a)**. Given the complimentary effect of the two antibodies, we infer that a combination of mAbs c6C with 1G10 are better experimental tools for understanding subpopulation(s) of EV and propose such a strategy for future countermeasure development. By either experimentally releasing the CEV from cells infected with VacV strain WR, or using those inherently released by the VacV strain IHDJ, we were able to show that these particles are still capable of A33-specific neutralization. Although this strategy isn’t too novel and likely already utilized by the immune system during the generation of a polyclonal response(56) combinations of multiple A33-based antibodies are rarely proposed, but could act to supplement already proposed combinations by acting as a “one-two punch” to both release and clear the CEV-like and EEV-like particles, in conjunction with other neutralizing mAbs specific for diverse EV protein targets (e.g., B5)(43, 50, 57). This would provide a more comprehensive approach for optimizing efficacy and possibly reducing tissue pathology. Our studies focused on functionality (productive infection), therefore we cannot surmise what effect, if any, mAb c6C has on the A33 structure or quantity within the virions, but these as well as other aspects will be addressed in future experiments. We can minimally conclude that in conjunction with the reported role of A33 and CEV release (58, 59{Chan, 2010 #1050, 60)}, that the activity of mAb c6C involves some non-A33 encoded virus-virus or virus-host interaction. This was further confirmed with our experiments involving the same virus in different cell lines (**Figure 5**) discussed later.

Although initially too risky to conduct the experiments needed to complement mAb c6C, our findings should not dissuade further testing of the mAb 10F10. Mabs 10F10 and 1G10 target specific epitopes present in most strains of poxviruses and the discrepancy seems solely based on the A33 sequence(49). This suggests that the breadth of protection against known or emerging A33 ortholog variants could be ensured by incorporating such a combination as part of a robust countermeasure.

Serendipitously, we found that we could cause aberrations concomitantly in both the quantity and mAb c6C sensitivity of VacV strain WR EV by only changing host cell type. That is without a change in the sequence of A33, we could change rCEV content; more specifically, by utilizing Vero 76 cells instead of HEK 293T cell lines (**Figure 5 A, B, and C**). These data were counterintuitive to our relatively modest reported quantities of EV produced by clade Ia MPXV in Vero E6 cells (45), but the comparison utilizing the exact same conditions and cell preparations allowed a wider perspective of possibilities for potential levels controlling EV (**Figure 5 and Figures S3 and S4**). Although overall replication kinetics may play a role between cell lines, for which we only had some anecdotal evidence (**Figure 6**), our interests were focused more on the specific composition or derivation of the population of EV based on CEV-like or EEV-like tendencies to neutralize (**Figure 5**). Although, our data shows that the exact trend seems specific to virus/virus strain, the major implication that similar mechanism(s) could be occurring during pathogenesis and transmission, between hosts and/or within the same host in different tissues/cell types, provides a broader avenue by which to understand poxviruses. This became more convincing by expanding our experiments with different MPXV strains. Although these studies were somewhat hampered by incomplete neutralization by any of our anti-MV or anti-EV mAbs typically utilized to confirm the quality of total EV-like particles. Therefore, we tended to have greater assay variability and relied on the overall common trend in total and rCEV-like populations (**Figure 5F**). Defining statically significant MPXV-strain specific differences in the future would require high repetitions of experiments or more sensitive measures of infectious EV populations.

Given the differences discussed thus far, we also wanted to understand how changes could relate to stability of the EV particle and determine the impact in virus transmission and/or intra-tissue/intra-host dissemination. We had previously shown that clade Ia MPXV EV propagated from NHP cells were sensitive to the effects of NHP complement (45) and unlikely to have a similar mechanism of resistance as previously reported for VacV EV; a mechanism by which host specific cellular proteins incorporated into the EV was primarily important for such species specific resiliency (44). Therefore, we further tested sensitivity to host specific complement utilizing clade Ia or IIb MPXV EV propagated in human or NHP derived cell lines (**Figure 1 C and D**) and compared the results to VacV-based experiments (**Figure 2**). The relative stability of freshly released MPXV clade I EV tended to be low for MPXV given that either anti-MV antibody or complement could decrease spread (45), therefore the effects were compared back to a known quantity of EV under anti-MV neutralizing conditions and disregarded the initial neutralization caused by EV-assay experimental conditions (**Figure 4 C, D and Figure 5**). Data reported here suggest that the cell type (host encoded protein) is less important than a virally encoded factor. Anecdotally, this can be confirmed with known genetic similarities between experimental VacV strains and differences between clade I and II of MPXV (e.g., lack of virally encoded complement control protein in clade II MPXV)(61). We were able to confirm findings noted by Vanderplasschen, et al with VacV strains being resistant to human sera and not rabbit sera based on EV derivation from human cells (44), but our findings do not extend to the proposition of the role of a cell line when we utilized Vero-cell line to propagate and test EV. Vero 76 derived EV had similar sensitivities to NHP complement and human complement, but not rabbit complement (**Figure 5**). Together, it appears the resilience of VacV EV to complement/sera may be broader to include both human and non-primate origin, but the resilience seems unrelated to the production cell type itself. Studies utilizing an array of assays of host-based cell lines with supportive molecular findings is necessary to precisely answer such a complex question. Our initial evidence suggests that under similar experimental conditions there are differences between MPXV and VacV, as well as between clades of MPXV, and they are likely virally encoded, but we cannot state a definitive nil effect of cellular incorporated proteins. Furthermore, the incorporation of host complement decay factors in VacV derived vectors for oncolytic treatment seems to provide more stability *in vivo* (62)

Poxviruses are very complex, with over 200+ encoded proteins and producing multiple infectious forms. The 2022 MPOX global outbreak by a new subclade of MPXV (IIb) and the continued endemic of the clade Ia and Ib viruses in Africa has shown that the threat of emerging poxviruses, our understanding of their virulence and transmission, and our ability to successfully predict, prevent, and/or treat a more novel poxvirus may be lacking. The MPOX scenario becomes more of a concern when compounded with our very limited understanding of smallpox and/or variola-like poxviruses. As we have just shown, studying characteristics and features of poxviruses *beyond* prototype viruses and utilized *common in vitro* and *in vivo* hosts is needed to increase our breadth of understanding. Nuances created by small, sometimes seemingly nominal, or even major changes in viral sequence can have different impacts at various stages of the viral cycle that can be missed due to the artificial and typically repetitive systems scientists tend to utilize. In the studies presented here, we focus on infectious EV-like particles in a very limited number of well characterized cell lines. We have yet to understand the effect of cell changes on viral intermediates, MV, or secreted viral factors that likely play a role in pathogenesis and transmission. Therefore, an overabundance of caution and continued study is always warranted to ensure protection from known, emerging, and other viral threats.

## Materials and Methods

### Cells, viruses, and antibodies

Briefly, cells were infected with either VacV WR, VacV IHDJ, MPXV strain USA 2004, and MPXV strain Zaire 76 were obtained and propagated as previously described(39, 63). EV neutralization assays utilized human complement (Cedarline and Bioreclamation) and anti-L1 antibody, 7D11 (USAMRIID). Antibody targeting vaccinia virus protein L1 (Mab 7D11), A33 (Mabs 1G10 and 10F10, USAMRIID; Mab c6C, BioFactura, LLC) and B5 (Mab c8A, BioFactura, LLC) have been described elsewhere(49, 50). Rabbit and cynomolgus macaque complement (BioReclamation) were utilized for stability assays.

### EV Neutralization

EV neutralization has been described elsewhere for VacV(**39**) and MPXV(45). For VacV strains, EV content was presented as a titration and/or percent of infectious virus remaining after neutralization with an anti-L1 antibody. Mucker et al. 2019 had shown that monoclonal antibody (Mab) against A33, known as Mab c6C, could reduce plaque size, neutralize VacV EEV, and induce, but not neutralize, the release of VacV CEV (rCEV) from cells. Therefore, Mab c6C neutralizable content are identified as more EEV-like, whereas the fraction not susceptible to Mab c6C are considered more CEV-like or rCEV-like. Content of VacV EV within the stocks were confirmed by use of anti-B5 antibody and shown to coincide with MV-based titers. For MPXV, total EV are presented as a titration of the stocks (PFU/mL) in the presence of a final concentration of 1:50 and 5% of Mab 7D11 and human complement, respectively. All neutralization calculations for anti-A33 and/or B5-antibodies were performed as a percent neutralization of a target of 100 PFU/well of EV that were empirically calculated from stocks after incubation with anti-L1 and human complement.

### CEV release assays

Pilot studies to determine conditions, impact Mab c6c concentration and timing of harvest on release of cell associated EV were also performed using BSC-1 cells seeded on 6 well plates (**Figure S3 B**). Varying concentrations of Mab c6C were added to monolayers of BSC-1 cells infected at an MOI of 1 of VacV as indicated. Supernatants were then collected after two days, clarified twice via centrifugation at 800 x g for 10 minutes, and EV neutralization assays were performed utilizing additional one of two Mabs, c6C or IG10, with 121.5 ug/mL or treated with an equal volume of media. Data were normalized to the mock for each Mab c6C concentration tested.

### Stability testing of VacV and MPXV EV stocks

For complement assays, stocks were diluted to target concentration of 100 PFU for each EV preparation and were incubated with either 5% or 50% human or cynomolgus macaque sera (Bio IVT) in the presence of anti-MV antibody (**Figure 1 and 2**). Controls utilizing matching heat inactivated sera at 50% were concomitantly performed to assess total EV/infectious virus content. Percent neutralization was calculated normalized to input EV. Stability over time was also tested in multiple EV preparations (minimum of N=3 per virus strain) and both titration of EV and %EV were calculated at the timepoints provided in the figures.

### In vivo samples and processing

All samples were run (>2 week) without freeze thaws or sonication. Samples from two separate, but ongoing studies utilizing cynomolgus macaques of Asian (Study 1) or Mauritius (Study 2) origins were collected and subjected to neutralization assays as indicated (**Table 1**). To limit the possibility of cells contributing to our results, serum was utilized. Oropharyngeal swabs were placed in EMEM containing antibiotics/antimycotics and 2% heat inactivated fetal bovine serum. Additional neutralization assays were performed concomitantly for Study 2 samples: samples were incubated in the presence of 5% human complement alone, 7D11 alone, or 5% complement and anti-EV Mab c8A in an effort to compare EV content by standard indirect neutralization (i.e., anti-MV antibody/complement and determine remaining infectious virus) or direct neutralization of the EV particle (5% complement and Mab c8A and identify % neutralized as EV population), as well as the contribution(s) to neutralization each component provides to the infectious nature of each sample.

### Electron microscopy

Cells were fixed with phosphate buffered 4% paraformaldehyde for at least 1 hour at room temperature then incubated overnight at 4C. After warming to room temperature, samples were rinsed twice with purified EM grade water then post-fixed with 1% osmium tetroxide in water for 45 minutes at room temperature. Samples were rinsed once with water and then scraped into pellets. Centrifuged pellets of cells were incubated with 1% ethanolic uranyl acetate for 30 minutes for additional contrast and nucleic acid fixation. Samples were then subjected to increasing concentration of ethanol for water extraction and dehydration. Following the 100% ethanol dehydration, samples were incubated with 100% ethanol and 100% propylene oxide [P.O.; 1:1] for 5 minutes before dehydrating with 100% P.O. Next, samples were incubated with P.O. and epoxy resin [2:1; EM sciences,] for 30 minutes and then P.O. and epoxy resin [1:1] for 30 minutes to overnight at room temperature. Next day, samples were incubated and subjected to three changes with 100% resin. Labels were inserted in each sample and allowed to polymerize overnight at 60°C.

#### Molecular viremia

Viral load was determined using a real-time RT-PCR assay targeting the MPXV F3L gene as previously described (64). Briefly, total nucleic acid was extracted from 100 μl sample by adding to 100 μl buffer ATL (Qiagen), incubating at 56°C for 15 min at 300 RPM, and processed using the EZ1&2 virus mini kit 2.0 (Qiagen). A standard curve was generated using the virus challenge stock. Viral load was determined by conducting real-time RT-PCR on five μl extracted nucleic acid using the TaqPath 1-Step RTqPCR Master Mix, CG kit (ThermoFisher Scientific) according to the manufacturer’s instructions. Primers MPOX-F3L-F290, MPOX-F3L-R396a, and probe MPOX-F3L-p333 were sed on the QuantStudio Dx (ThermoFisher Scientific) using the following cycling conditions: 25°C x 2 min; 50°C x 15 min; 95°C x 2 min; 45 cycles (95°C x 3 sec, 60°C x 30 sec). A sample was called positive is the Cq value was below 40 cycles, and the viral load was determined based on the standard curve.

### Antigen assays and serology (Magpix)

The presence of mpox antigen in serum was determined with a sandwich-style immunoassay previously developed on the Magpix platform(65).Briefly, sera were diluted 1:20 in PBS with 5% skim milk powder and 0.05% Tween. 50 µL of diluted sera were incubated with magnetic microspheres functionalized with a mpox specific antibody for 1 h. The microspheres were washed three times and incubated with a phycoerythrin (PE)-labeled secondary antibody for 1 h. The microspheres were then washed three times and read by the Magpix instrument. The median fluorescent intensity (MFI) from the PE reporter is proportional to the amount of antigen present in the sample.

Serology was performed with a multiplexed indirect immunoassay previously developed on the Magpix platform (66)Briefly, sera were diluted 1:100 in PBS with 5% skim milk powder and 0.05% Tween. 50 µL of diluted sera were incubated with antigen-functionalized, barcoded magnetic microspheres for 1 h. A different barcode was used for each antigen. The antigens used were MPXV A29, A35, B6, and VACV A27, A33, B5, and L1. The microspheres were washed three times and incubated either anti-human IgG-PE or anti-human IgM conjugate for 1 h. The microspheres were washed three times and read by the Magpix instrument. The data is reported as MFI divided by the MFI from day 0 for each NHP.

### Statistics

All descriptive statistics were performed utilizing GraphPad Prism Version 9.4 software.

## Disclaimer

The opinions, interpretations, conclusions, and recommendations presented are those of the author and are not necessarily endorsed by the U.S. Army or Department of Defense.

The use of either trade or manufacturers’ names in this report does not constitute an official endorsement of any commercial products. This report may not be cited for purposes of advertisement

USAMRIID is fully accredited by the AAALAC International, and all animal research conducted at USAMRIID complies with the Animal Welfare Act and other federal statutes and regulations relating to animals in research and adheres to The Guide for the Care and Use of Laboratory Animals, National Research Council, 2011.

